# Genetic determinants of intrinsic antibiotic tolerance in *Mycobacterium avium*

**DOI:** 10.1101/2021.02.23.432616

**Authors:** William M. Matern, Harley Parker, Carina Danchik, Leah Hoover, Joel S. Bader, Petros C. Karakousis

## Abstract

*Mycobacterium avium* complex (MAC) is one of the most prevalent causes of nontuberculous mycobacteria pulmonary infection in the United States, yet it remains understudied. Current MAC treatment requires more than a year of intermittent to daily combination antibiotic therapy, depending on disease severity. In order to shorten and simplify curative regimens, it is important to identify the innate bacterial factors contributing to reduced antibiotic susceptibility, namely antibiotic tolerance genes. In this study, we performed a genome-wide transposon screen to elucidate *M. avium* genes that play a role in the bacterium’s tolerance to first- and second-line antibiotics. We identified a total of 193 unique *M. avium* mutants with significantly altered susceptibility to at least one of the four clinically used antibiotics we tested, including two mutants (in DFS55_00905 and DFS55_12730) with panhypersusceptibility. The products of the antibiotic tolerance genes we have identified may represent novel targets for future drug development studies aimed at shortening the duration of therapy for MAC infections.

**Importance:** The prolonged treatment required to eradicate *Mycobacterium avium* complex (MAC) infection is likely due to the presence of subpopulations of antibiotic-tolerant bacteria with reduced susceptibility to currently available drugs. However, little is known about the genes and pathways responsible for antibiotic tolerance in MAC. In this study, we performed a forward genetic screen to identify *M. avium* antibiotic tolerance genes, whose products may represent attractive targets for the development of novel adjunctive drugs capable of shortening curative treatment for MAC infections.

## Introduction

Nontuberculous mycobacteria (NTM) are found ubiquitously in the environment and several species can cause disease especially in the elderly, those with preexisting lung disease, and the immunocompromised, including those infected with HIV (1–4). *Mycobacterium avium* complex (MAC), a group of 12 related, slow-growing mycobacteria with *M. intracellulare* and *M. avium* as the most prevalent species, accounts for the majority of pulmonary infections due to NTM in the US (5,6). Although the true incidence of pulmonary MAC infections in the US is not known, a study in Oregon reported 4.8 cases per 100,000 person-years in 2012 (7). Winthrop *et al.* estimated the annual incidence of NTM infections in the US to be 4.73 cases per 100,000 person-years (8), and the NTM-NET group has reported that MAC accounts for 52% of NTM isolates in the USA and Canada (9), suggesting the annual incidence of MAC in the US may be closer to ~2.5 per 100,000 person-years in the US.

Current treatment for MAC comprises a combination of multiple antibiotics given for at least 12 months following the conversion of sputum cultures from positive to negative. Since sputum culture conversion occurs in ~50% of cases after five months of antibiotic therapy, a typical patient receives a minimum of 15-18 months of treatment (10,11). Macrolide-susceptible MAC is usually treated with at least three antibiotics, including a macrolide (azithromycin or clarithromycin), a rifamycin (rifampin or rifabutin), and ethambutol, either intermittently (three times weekly) or daily for severe fibronodular or cavitary disease. If the infecting MAC strain is macrolide-resistant or the patient is unable to take the first-line regimen, alternative antibiotics, such as moxifloxacin, clofazimine, or linezolid are often used (11,12).

The lengthy and complicated treatment course required to eradicate MAC infection has been attributed to the presence of persistent organisms, which exhibit reduced susceptibility, or tolerance, to antibiotics (13). Unlike antibiotic resistance, which results from a heritable genetic alteration permitting continued bacterial growth in the presence of antibiotic concentrations exceeding the minimum inhibitory concentration (MIC), antibiotic tolerance is a transient, nonheritable phenotype without associated change in the MIC. The term “antibiotic tolerance” was originally coined in 1970 by Tomasz *et al.* to denote the ability of bacteria to withstand the bactericidal activity of antibiotics, especially of cell wall-active agents, primarily by reducing their replication rate (14). In the intervening decades, additional mechanisms have been proposed to mediate bacterial antibiotic tolerance, including biofilm formation (15,16), the stringent response (17–20), induction of efflux pumps (21–23), and altered metabolism (24,25). Following ingestion by macrophages, MAC acquires an antibiotic-tolerant phenotype within the arrested phagosome (26). Moreover, various stress conditions, including nutrient starvation, low pH and hypoxia, induce a nonreplicative, antibiotic-tolerant state (24), which is characterized by transcriptional changes (24,25), leading to altered cell wall membrane permeability (3) and increased expression of efflux pumps (21). This stress-induced adaptation of MAC is accompanied by dramatically reduced metabolism, with a shift to the glyoxylate shunt, stabilization of the mycolate pool, and a switch to transcription of only essential genes (25). Additionally, the glyoxylate shunt enzyme isocitrate lyase is critical for long-term survival of the related pathogen *M. tuberculosis* in host tissues (27), and can be used in a reductive amination pathway to produce NAD, which may serve as an alternative energy source in a nonreplicative state and under anaerobic conditions (28,29). However, the molecular mechanisms driving antibiotic tolerance in MAC remain poorly understood.

Transposon insertion sequencing (Tn-seq) is a powerful technique for determining bacterial genotype-phenotype relationships, particularly the requirement for specific bacterial genes for growth and/or survival under controlled stress conditions (30–33). Modifications of this technique have been used to define essential genes for *in vitro* growth of *M. tuberculosis* (34) and *M. avium* (35). Xu *et al.* screened a saturated transposon mutant library in the presence of partially inhibitory concentrations of various antibiotics with diverse mechanisms of action to identify genetic determinants of intrinsic antibiotic susceptibility in *M. tuberculosis* (36). In the current study, we used a similar approach to identify genes responsible for intrinsic tolerance of *M. avium* to the antibiotics clarithromycin (CLR), rifabutin (RFB), moxifloxacin (MOX), and ethambutol (EMB). The hits we have identified may serve as targets for the development of novel antibiotics, with the objective of shortening the duration of curative treatment for MAC.

## Methods

### Strains

All experiments were performed using *Mycobacterium avium* subsp. *hominissuis* strain 109 (35).

### Media and buffers

To make 7H11 agar, 10.25 g of 7H11 w/o Malachite Green powder (HiMedia Cat No. 511A) was added to 450 mL deionized water. A volume of 5 mL 50% glycerol was then added before autoclaving. The agar was cooled to 55° C before addition of 50 mL OADC enrichment (Becton Dickinson). To make 7H9/30% OADC, 2.35 g of 7H9 powder was added to 350 mL deionized water. After sterilization (by autoclaving at 121° C or passing through a 0.22 μm filter) 150 mL of OADC enrichment was added. Unless otherwise specified, no Tween-80 or glycerol was included in the media. To make PBS-Tw, 1.25mL filter-sterilized 20% Tween-80 was added to 500 mL of sterile phosphate-buffered saline (PBS).

### Overview of transposon screen setup

The setup for our genome-wide differential susceptibility screen is shown in Figure 1. We inoculated a 1-mL frozen aliquot of a previously generated diverse *Himar1* transposon mutant pool (consisting of approximately 1.2 × 10^6^ unique mutants) (35) into 300 mL of 7H9/30% OADC contained in a 1.3-L roller bottle. This was shaken at 37°C for 24 hours to reduce bacterial clumping (220 rpm, 0.75 in [1.905 cm] orbit). The optical density at 600 nm (OD_600_) was tracked until exceeding 0.32. The transposon mutants were then diluted to OD_600_ 0.1 with cold 7H9/30%OADC and aliquoted into 89 50-mL conical tubes (10 mL/tube). Tubes were incubated for 5 hours while shaking at 37° C to allow the bacteria to return to log-phase growth. Five tubes were randomly selected for processing (0 hours). These samples were then processed for colony forming-unit (CFU) enumeration and regrowth (same tube used for both operations). Following regrowth, bacterial samples were scraped and processed for Tn-seq. Regrowth is required before Tn-seq library preparation in order to remove the contribution of DNA to the sequencing library from dead transposon mutants. After collection of enumeration and regrowth samples at 0 hours, antibiotics (or DMSO vehicle as control) were then added to all tubes. Samples were collected at 12 and 48 hours after antibiotic exposure in triplicate and processed for CFU enumeration and regrowth in the same manner as the 0 hours samples.

**Figure 1:**
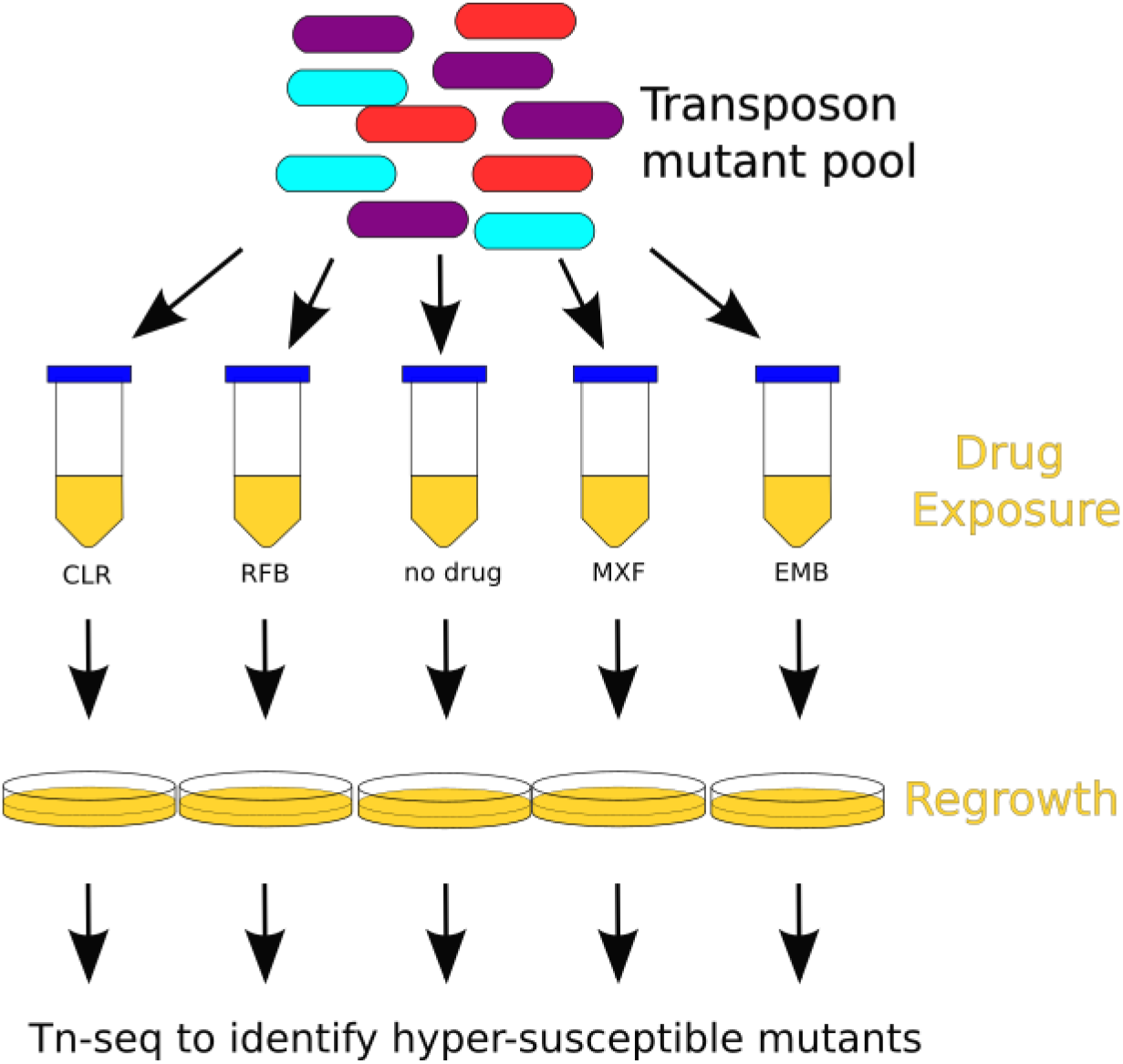
Schematic of transposon mutant screen to identify hypersusceptible mutants following exposure to multiple doses of various antibiotics in liquid culture. After antibiotic exposure, cultures were regrown on solid agar to enrich for surviving bacteria. After regrowth, DNA was extracted and prepared for Tn-seq analysis. Hypersusceptible mutants were identified using a non-parametric statistical approach.

### Sample processing for CFU enumeration and regrowth

The bacterial density (CFU/mL) was estimated by removing 400 μL of bacterial culture, centrifuging (2000 g for 5 minutes) and washing twice with PBS-Tw to remove antibiotic. Washed samples were diluted 10-fold and 50 μL of each dilution was plated on 7H11 agar without malachite green. Malachite green, which is typically present in standard 7H10 and 7H11 agar formulations, was excluded to avoid potential issues with post-antibiotic recovery in mycobacteria (37). T-shaped spreaders were used to spread liquid evenly across agar plates. CFU were counted after 7-8 days.

For regrowth, the remainder of each tube (after removing samples for CFU enumeration) was centrifuged twice and washed (2000 g for 10 min) with 10 mL of PBS-Tw to remove antibiotic. The samples were centrifuged once more and the bacterial pellet was resuspended in 250 μL PBS-Tw. Fifty μL of the washed transposon pool was plated on each of four 7H11 agar plates and spread with 10-15 3-mm sterile glass beads to ensure even distribution of liquid across the plate. Samples were regrown for 7-8 days. Bacterial lawns from the four agar plates were scraped and pooled into 2-mL tubes. DNA was extracted from regrown samples, as described previously (short-read sequencing protocol) (38). DNA was processed for Tn-Seq, as described previously (35). Libraries were sequenced (2×75bp) on an Illumina HiSeq 2500 by the Johns Hopkins GRCF High Throughput Sequencing Center. A total of 59 samples (5 input-pool samples and 18 groups of output-pool triplicates) were sequenced, yielding between 2,333,295 – 7,193,522 reads per sample for a total of 269,324,560 paired-end reads.

### Antibiotic selection

Doses were selected to reflect antibiotic concentrations at the most common site of infection in non-compromised patients (the lungs) following standard antibiotic dosing. Based on a search of the pharmacokinetic literature, maximum achievable doses in lung tissues were taken to be 54 μg/mL for CLR (39), 0.63 μg/mL for RFB (40), 10.0 μg/mL for MOX (41), and 21.0 μg/mL for EMB (42) (based on non-human primate data). For each drug, a 10-fold dilution series of the estimated maximum achievable dose was performed to explore the impact of dose. A preliminary calibration experiment was performed to estimate the bacterial viability at different antibiotic concentrations (data not shown) and to select the number of concentrations to include (4 total concentrations for CLR, 3 concentrations for the other antibiotics in addition to drug-free controls).

### Sample selection for sequencing from susceptibility screen

Given resource constraints, only a subset of samples from the differential susceptibility screen was chosen for Tn-seq library prep and sequencing. Samples were sequenced at two manually chosen concentrations for each drug at both available timepoints (12 hrs, 48 hrs). To help identify which should be processed further, objective criteria were established a priori, with the goal of clearly identifying mutants with higher susceptibility to antibiotics, but also reducing the likelihood that mutants were removed by chance due to low bacterial viability during antibiotic exposure. We considered the following 3 criteria for selecting samples:

A. Bacterial numbers must exceed 10^6^ CFU/mL at all times during exposure. This ensures that the probability of losing a non-defective mutant is minimized, given the 60,129 possible thymine-adenine (TA) sites for the *Himar1* transposon to insert across the MAC109 genome. Only CFU data were used, and we did not attempt to directly estimate the total number of bacterial cells, which can differ significantly from CFU.
B. There should be decreased bacterial viability after antibiotic exposure relative to drug-free controls (as measured by CFU). This ensures the concentration of antibiotic is high enough to have bactericidal activity. Otherwise, the antibiotic concentration might be too low to select for or against mutants with growth phenotypes.
C. Drug concentrations near or below achievable serum concentrations of the drug after standard dosing are preferred. We assumed approximate serum values of 2.31 μg/mL for CLR, 4.42 μg/mL for MOX, 0.52 μg/mL RFB and 2.27 μg/mL for EMB (43). In our view, this criterion makes the results more clinically relevant.

### Identification of hypersusceptibile mutants from sequencing data

A schematic of the pipeline to process the data is provided in Figure S1. Raw reads were mapped using the TRANSIT preprocessor (tpp) (44). Counts from tpp were then processed with a custom python script to produce a *.csv file to be read by pandas (version 0.24.1) used for downstream analyses.

### Effect size/log-fold change calculation

For calculation of the normalized read counts for each mutant, a pseudocount of 4 was added to the raw count from all samples (for stabilization) before dividing read counts by total read count:

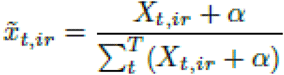

(where *x_t,ir_* is the normalized read count, *X_t,ir_* is the raw read count for transposon insertion site *t*, for antibiotic treatment group *i*, for replicate *r*. *α* is the pseudocount (α = 4) and *T* is the number of transposon insertion sites). This pseudocount was determined by manual examination of read counts in genes known to be essential and set to be substantially larger than occasional background reads. The normalized read counts were then averaged over samples:

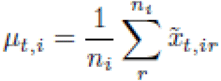

(where *μ_t,i_* is the average representation of each mutant across samples, and *n_i_* is the number of replicates for treatment group *i*). An aggregate log-fold-change (LFC) was used as a measure of effect size for differentially susceptible mutants. The aggregate LFC between treatment groups *i* and *j* was calculated as the median of the log-fold change change at individual transposon insertion sites within a gene:

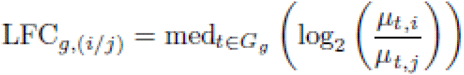

where *G_g_* is the set of transposon insertion sites annotated to belong to gene *g* and *LFC_g,(i/j)_* is the log-fold change between treatment groups *i* and *j* for gene *g*. Although this formula is generally useful for comparing any pair of groups, for the work presented here, *i* always represents a drug-containing treatment group and *j* always represents the matching drug-free group at the same time point

### P-value calculation

For calculation of p-values, read counts for each sample were first normalized by dividing by the total read count in each sample (pseudocounts are unnecessary for the non-parametric test described below, and thus not used).

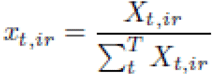

The Jonckheere-Terpstra (JT) test was then applied to the normalized read counts at each time point (45,46). Briefly, the JT test is a non-parametric test of trend, which is more powerful than the more commonly used Kruskal-Wallis test when the alternative hypothesis assumes a monotonic trend of the treatment groups. In this case, we have three treatment groups for each drug at each time point: No drug, low dose, high dose. We postulated that if a mutant is hypersusceptible at a low dose of antibiotic, it will be even more hypersusceptible at a higher dose of that antibiotic. We define the alternative hypothesis as:

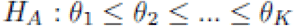

against a null hypothesis:

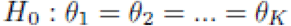

(where *θ_1_...θ_K_* are measures of a centrality parameter and *K* is the number of treatment groups for the drug at a particular time point (in this case *K* = 3 for each drug)). The JT test statistic (B_t_) at site *t* is defined as:

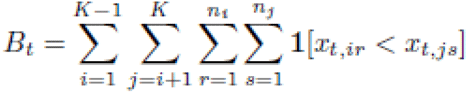

(where **1[ ]** is the indicator function). P-values at individual sites are computed via permutation test. Naively, this would result in non-uniformly distributed p-values due to the discrete nature of the distribution. To ensure a truly uniform distribution, we performed a small correction on the permutation test p-values by sampling from a uniform distribution bounded between adjacent values of the discrete permutation distribution. This process gives precisely uniform p-values under the null hypothesis.

Pooling the p-values within each gene (one p-value calculated for each transposon insertion site) is then accomplished with the two-sided Stouffer’s method. Finally, adjusted p-values are then computed using the Benjamini-Hochberg procedure.

### Thresholds for defining differentially susceptible mutants

Gene mutants are considered “differentially susceptible” to a drug if the absolute value of the LFC at the high dose of each drug (relative to no drug control) is greater than 0.5 and the adjusted p-value is less than 0.05. This condition must be met at both the 12 hours and 48 hours time points in order for the mutant to be defined as differentially susceptible to the drug. Mutants with negative LFC are predicted to be hypersusceptible to the drug, while positive LFC indicates the mutant is hypertolerant.

### Raw data and code availability

The raw sequencing data (*.fastq) from this project can be found in the NCBI Sequence Read Archive (SRA) under BioProject number PRJNA559896. A Jupyter notebook and associated scripts to reproduce the data analysis (including figures) from the raw data is provided on github.com (https://doi.org/10.5281/zenodo.4542412).

## Results

### Effects of antibiotics on bacterial growth at the population level

To monitor the effects of individual antibiotics on the entire bacterial population, we measured CFU and OD_600_ during antibiotic exposure. CFU values obtained following antibiotic exposure for 0, 12, and 48 hrs are provided in Figure S2. OD_600_ values are provided in Figure S3. The same no-drug (vehicle) control data appear in all four plots (performed in triplicate). Notably, the no-drug control curve has an inflection at the 12-hr time point. Truly logarithmic growth should appear as a straight line on this plot. Light microscopy of unstained samples of the no-drug control cultures revealed clumps of approximately 5 bacteria (data not shown), likely accounting for the inflection point. Bacterial clumping was likely also present in the antibiotic-containing tubes, although these were not specifically examined. The presence of clumping is unlikely to impact the results of the screen, as there is no reason to suspect that individual transposon mutants were disproportionately distributed among the clumps. Applying the set of criteria for selecting samples for processing, we prepared libraries and sequenced both time points at the following concentrations: CLR 0.54 and 5.4 μg/mL, MOX 0.1 and 1.0 μg/mL, RFB 0.063 and 0.63 μg/mL, EMB 2.1 and 0.21 μg/mL. The colored arrows in Figures S1 and S2 indicate the time points sequenced.

### Identification of mutants with altered antibiotic susceptibility

A total of 161 mutants showed increased susceptibility and 32 mutants showed reduced susceptibility to at least one of the four antibiotics. 46 mutants were hypersusceptible and 14 mutants were hypertolerant to CLR. 6 mutants were found to be hypersusceptible to EMB, while no mutants were hypertolerant to this antibiotic. The MOX screen revealed 103 hypersusceptible and 2 hypertolerant mutants. 84 mutants were found to be hypersusceptible and 108 mutants were hypertolerant to RFB. Effect sizes (after 48 hours of exposure) for mutants with significantly altered antibiotic susceptibility are plotted in Figure 2 and summary data are provided in Tables S3-S6.

**Figure 2:**
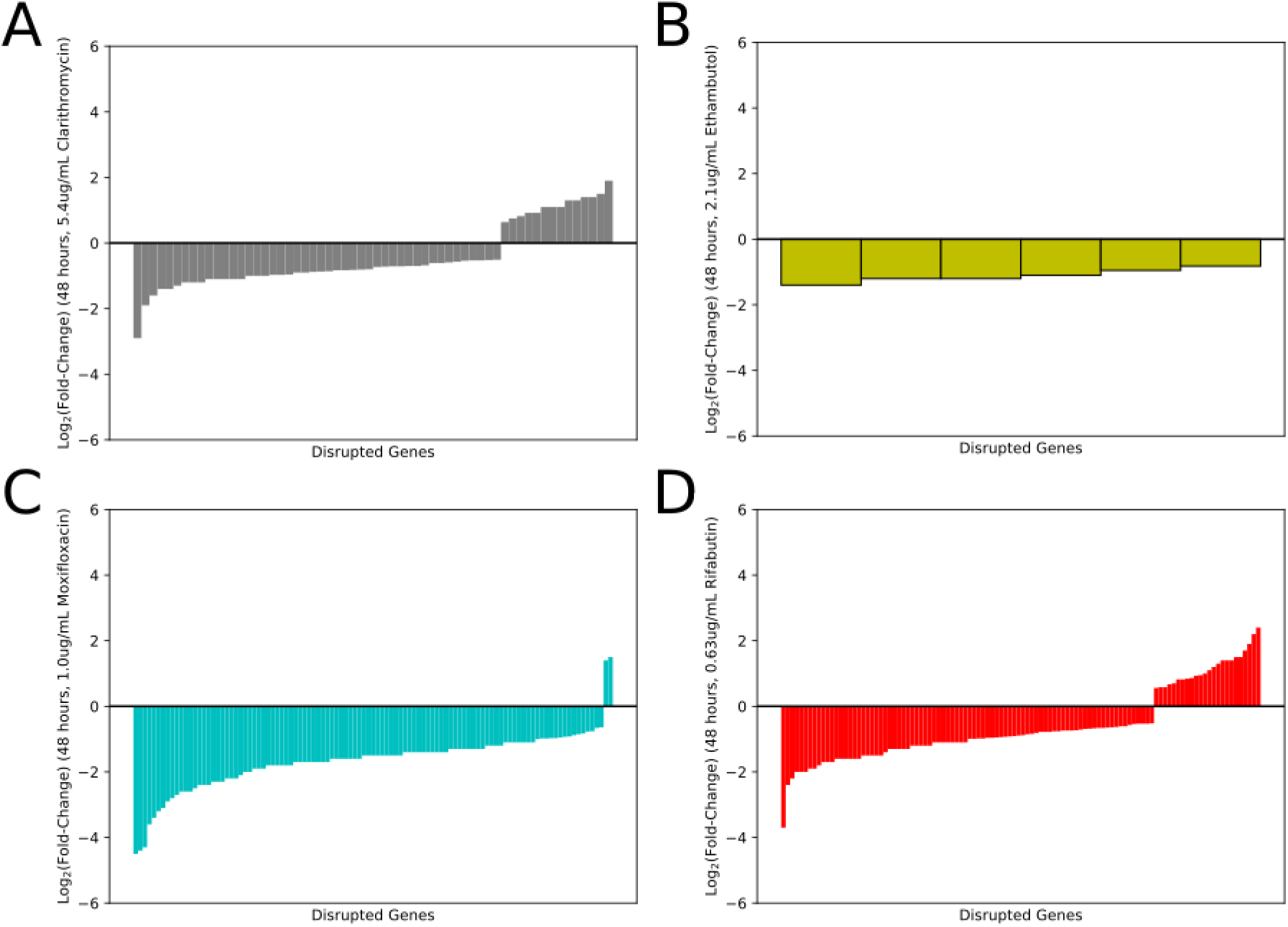
Bar chart showing the effect size of each statistically significant mutant. Each bar represents a single gene. A negative value represents a hypersusceptible mutant, while a positive value signifies that a mutant is less susceptible (hypertolerant) to the antibiotic. (A) clarithromycin, (B) ethambutol, (C) moxifloxacin, (D) rifabutin.

We also looked for overlaps between the different drug classes tested. Our results of this analysis are summarized in Figure 3 and Table S1 (hypersusceptible mutants) and Table S2 (hypertolerant mutants). Mutants hypersusceptible to multiple antibiotics may reflect genes with a role in more general bacterial persistence mechanisms, while mutants hypertolerant to multiple antibiotics may suggest genes promoting antibiotic susceptibility. Notably, no mutant was found to be hypersusceptible to one antibiotic but hypertolerant to another.

**Figure 3:**
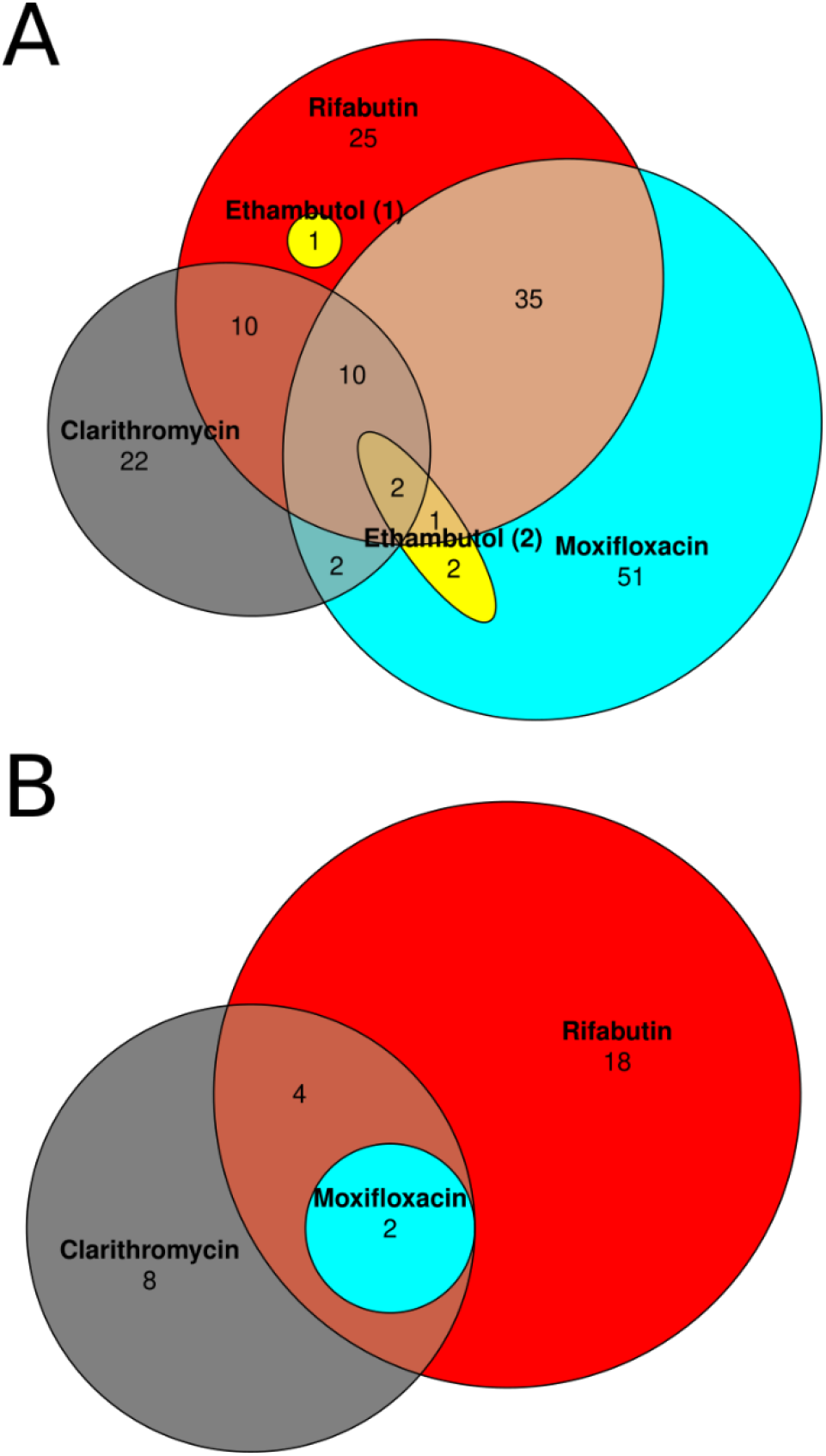
Venn diagram of identified hypersusceptible (Panel A) and hypertolerant transposon mutants (Panel B). Note that in Panel A the set of ethambutol-hypersusceptible mutants has been partitioned into two sets (both in yellow). Partitioning in this way greatly simplifies the diagram. Gene names in each category can be found in Tables S1-S2.

Two mutants were hypersusceptible to all four antibiotics tested, namely DFS55_00905 (annotated as an acyltransferase, homologous to *M. tuberculosis* Rv0111) and DFS55_12730 (hypothetical protein, homologous to Rv1836c). 10 mutants were hypersusceptible to CLR, MOX, and RFB, but not to EMB.

Two mutants were identified as hypertolerant to CLR, MOX, and RFB (no mutant was hypertolerant to EMB): DFS55_10765 (annotated as a pyruvate kinase, homologous to Rv1617) and DFS55_20040 (DUF1707 domain-containing protein, homologous to Rv0966c). An additional 4 mutants were found to be hypertolerant to RFB and CLR only, but not MOX. These included DFS55_10660 (quinolinate synthase, homologus to Rv1594), DFS55_10665 (L-aspartate oxidase, homologous to Rv1595), DFS55_16845 (trigger factor, homologous to Rv2462c), and DFS55_21750 (hypothetical protein, homologous to Rv3489).

## Discussion

In this work, we utilized a genome-wide transposon mutant pool to screen for *M. avium* mutants with altered susceptibility to various clinically relevant antibiotics. Compared to the other antibiotics, exposure to MOX yielded the highest number of hypersusceptible mutants, highlighting the many potential targets which might synergize with this antibiotic. Also, given the strong effect sizes observed with MOX relative to the other antibiotics tested, MOX synthetic lethality may represent the greatest opportunity for novel treatment-shortening strategies.

Transposon insertions in several known virulence genes were found to enhance susceptibility of *M. avium* to multiple antibiotics. For example, mutations in *secA2* (DFS55_12665 or *rv1821*) conferred hypersusceptibility to both CLR and MOX. The Sec export pathway is conserved across bacteria and exports secreted proteins across the cytoplasmic membrane (48). Mycobacteria have two SecA proteins, SecA1 and SecA2 (48). While SecA1 is essential and facilitates the transport of unfolded proteins though the SecYEG channel via its ATPase activity, the mechanism of export in the SecA2 pathway is less well understood (49). SecA2 is required for secretion of *M. tuberculosis* virulence proteins and arresting phagosome maturation by preventing acidification, thereby facilitating *M. tuberculosis* growth within macrophages (50). In particular, the SecA2-secreted phosphatase SapM and the kinase PknG have been identified as effectors with direct roles in preventing phagosome maturation and promoting *M. tuberculosis* intracellular survival and replication (51). As CLR inhibits protein synthesis (52) and SecA2 disruption impairs secretion of virulence-related proteins, these two alterations which both dysregulate proteostasis may have a synergistic or additive effect, leading to higher antibiotic susceptibility for this mutant. A similar but more indirect mechanism could be proposed for the sensitization of this mutant to MOX, which inhibits DNA gyrase and topoisomerase IV (52). Both of these enzymes are involved in the winding and unwinding of DNA and are necessary for DNA replication and RNA transcription (53,54). MOX-induced reductions in mRNA transcripts may also dysregulate proteostasis in MAC, potentially explaining the similar phenotypes observed with CLR exposure.

RecA (DFS55_08530 or Rv2737c), which was found to be required for MAC tolerance to both MOX and RFB, plays a critical role in the mycobacterial DNA damage response, specifically in the repair of double-stranded breaks, as *M. smegmatis* cells lacking RecA are more sensitive to DNA damage (55). Specifically, after RecBCD resects double-stranded breaks, RecA is loaded onto the 3’ end of the DNA, helping to mediate a homology search and subsequent strand invasion (55). By inhibiting DNA topoisomerases, MOX promotes DNA damage and triggers a mutagenic SOS response, which can lead to the formation of persister cells (56). RecA activation promotes the self-cleavage of LexA, which is able to downregulate the SOS regulon (57). In turn, this suppression of the SOS regulon decreases persister formation. Chemical inhibition of RecA with suramin in DNA gyrase-depleted cells has been shown to improve killing of *M. tuberculosis* by several anti-TB drugs, including rifampin and EMB (56). RFB, the rifamycin tested in our study, inhibits DNA-dependent RNA polymerase and suppresses RNA synthesis. Previous work showed that a *recA*-deficient *M. tuberculosis* mutant was unable to develop resistance to rifampin, possibly due to inability to generate an SOS response (58). In short, it appears that inhibiting RecA, thereby suppressing the SOS response, could provide a means to decrease persister formation and improve killing of pathogenic mycobacteria, such as *M. tuberculosis* and MAC.

Interestingly, we identified two mutants as hypersusceptible to all four antibiotics tested in our study (Figure 3). These included mutants with transposon insertions in DFS55_00905 (annotated as an acyltransferase, homologous to *rv0111*) and DFS55_12730 (hypothetical protein, homologous to *rv1836c*). Mutants in DFS55_00905 displayed particularly robust hypersusceptibility to MOX (effect size: −2.0 at 1 μg/mL and 48-hr exposure) and EMB (effect size: −1.4 at 2.1 μg/mL, 48 hrs, which was the largest effect size we observed with this drug at 48 hrs). Mutants in DFS55_12730 were strongly hypersusceptible to CLR (effect size:-1.6 at 5.4 μg/mL, 48 hrs), MOX (−1.9 at 1.0 μg/mL, 48 hrs), and RFB (−1.5 at 0.63 μg/mL, 48 hrs). Future work should investigate the function of these gene products and their relationship to the pansusceptibility phenotype observed.

An additional 10 mutants were found to be hypersusceptible to CLR, MOX, and RFB, but not to EMB. These included mutants in *sigE* (DFS55_18590, *rv1221*) and an alpha-beta hydrolase gene (DFS55_15065, homologous to *rv2224c*, also known as *caeA* or *hip1*). Deficiency of sigma factor E (SigE) has been shown to confer increased susceptibility of *M. tuberculosis* to multiple drugs, including EMB and rifampin, but not to ciprofloxacin (59). These results differ somewhat from the results of our study. Upon closer examination of our results for EMB, we find that this mutant was barely outside our conservative thresholds for defining hypersusceptible mutants. While adjusted p-values at both time points were below cutoff (0.045 and 0.0001, cutoff: 0.05), the corresponding log-fold-changes were barely above our chosen thresholds (−0.47 and −1, cutoff: −0.5). Therefore, it is possible that our stringent cutoffs misclassified this mutant as displaying similar EMB susceptibility as wild type. On the other hand, the discrepancy regarding hypersusceptibility of *sigE*-deficient mycobacteria to fluoroquinolones may be due to differences between the two species (*M. tuberculosis* vs. MAC, which have vastly different growth rates) and/or the experimental designs (resazurin microtiter assay vs. our Tn-seq screen). Additional studies are required to further evaluate the impact of *sigE* deficiency on MAC susceptibility to EMB and fluoroquinolones. Consistent with our data in MAC, *M. tuberculosis* mutants in *caeA/hip1/rv2224c* have been shown to be hypersusceptible to rifamycins (rifampin) and macrolides (erythromycin) (60). Our data suggest that *caeA* deficiency also confers enhanced susceptibility to fluoroquinolones, which might be useful for designing novel therapies for *M. tuberculosis*.

We found 2 mutants that displayed hypertolerance to three of the four antibiotics tested (CLR, MOX, RFB): DFS55_10765 (annotated as a pyruvate kinase, *rv1617*) and DFS55_20040 (DUF1707 domain-containing protein, *rv0966c*). Interestingly, Rv1617 deficiency is associated with a large growth defect in *M. tuberculosis* (34,35), but the same phenotype is not observed in DFS55_10765-deficient *M. avium* (35). This suggests that the metabolic impact of pyruvate kinase deficiency is remarkably different between MAC and *M. tuberculosis*. Pyruvate kinase catalyzes the transfer of a phosphate group from phosphoenolpyruvate to ADP (yielding pyruvate and ATP). Central metabolism may be disrupted in bacteria lacking this enzyme, possibly triggering the stringent response, which has been previously shown to protect bacteria from antibiotic-mediated killing (61,62). Deletion of pyruvate kinase in *M. tuberculosis* causes allosteric inhibition of the TCA cycle through accumulation of phosphoenolpyruvate (63). Disruption of the TCA cycle, especially alternate catabolism through the glyoxylate shunt, has been linked to antibiotic tolerance in multiple bacterial species, including *P. aeruginosa* (64), *S. aureus* (65,66), and *S. epidermis* (67), suggesting that this pathway may be a common mechanism for promoting antibiotic tolerance. In *M. tuberculosis*, downregulation of the malate synthase GlcB, one of the enzymes in the glyoxylate shunt, caused increased susceptibility to both rifampin and to nitrosative and oxidative stresses *in vitro* (68). Deficiency of isocitrate lyase, another glyoxylate shunt enzyme, also led to increased susceptibility of *M. tuberculosis* to antibiotics *in vitro* (69) and decreased survival in activated macrophages and mice (27). Thus, mycobacterial metabolism, and the TCA cycle in particular, clearly plays an important role in the development of antibiotic tolerance, although more work is necessary to fully elucidate its contributions. DFS55_20040 appears to lack an annotated function in the literature. Additional work is needed to understand the function of these two genes and determine their relationship to antibiotic tolerance in mycobacteria.

Our approach has several limitations. First, mutants in essential genes (or those without TA insertion sites) could not be screened, as these could not be recovered with our regrowth techniques. Therefore, we are unable to assess the potential role in antibiotic tolerance of genes essential for growth of *M. avium* in nutrient-rich medium. Second, gene disruptions leading to changes in secreted factors (e.g., extracellular proteins) may be missed by our screen, as these factors may be complemented by factors produced by non-defective mutants present in the same culture. Lastly, we chose a conservative statistical approach (JT-test) and conservative thresholds for p-values and LFCs, which must be met at two different time points. It is likely that mutants with low numbers of insertion sites or somewhat weaker effect sizes were missed.

Previous studies have examined mycobacterial antibiotic hypersusceptibility in the context of very low antibiotic concentrations (36). In such an experimental setup, the entire bacterial population continues to grow during antibiotic exposure and libraries were generated directly from bacterial cultures. In contrast, our approach here uses an additional regrowth step on solid agar after antibiotic exposure. This regrowth step produces sufficient material for library generation independent of whether the aggregate bacterial population is growing, stable, or dying. Therefore, our approach is more generally applicable to clinical scenarios in which higher doses of antibiotics may be used, entirely inhibiting aggregate mycobacterial growth.

Mutations causing defects while the aggregate population declines or is static are interpreted in our screen as amplifying the killing effect of the antibiotic (given that the wildtype organisms can be assumed to be non-growing). However, an observation of hypersusceptibility in the context of aggregate growth is more difficult to precisely resolve. Thus, it could be that the mutant is killed in the presence of antibiotic, whereas the wild type is able to grow, or it is possible that the mutant is more inhibited by the antibiotic relative to wild type, but continues to grow, albeit at a slower rate. In particular, the overall population declined in the presence of 5.4 μg/mL CLR, suggesting that any defective mutants are killed more rapidly than the wild type. However, at CLR 0.54 μg/mL, the overall population increased, suggesting that defective mutants could either be killed more rapidly or merely that their growth is inhibited to a greater extent than that of wild type (see Figure S2). Follow-up studies are needed in order to resolve the behavior of hypersusceptible mutants at this dose.

Our study represents a first step towards the development of novel, treatment-shortening strategies for MAC infections through identification of genes affecting antibiotic susceptibility. Biochemical characterizaation of the corresponding gene products might yield novel insights into the mechanisms of MAC antibiotic tolerance and lay the groundwork for the development of novel antibiotics, which might synergize with currently available drugs to more effectively kill tolerant organisms and shorten curative treatment for MAC infections. Future work is needed to validate the susceptibility phenotypes of individual gene-deficient mutants and their respective complemented strains in axenic culture. Proof-of-concept studies could then be performed to demonstrate the treatment-shortening potential of candidate targets in a relevant animal model of pulmonary MAC disease (70,71).

## Acknowledgements

This publication was made possible by a pilot research grant from the Sherrilyn and Ken Fisher Center for Environmental Infectious Diseases, Division of Infectious Diseases of the Johns Hopkins University School of Medicine. Its contents are solely the responsibility of the authors and do not necessarily represent the official view of the Fisher Center or Johns Hopkins University School of Medicine.

## Competing Interests Statement

WMM, HP, CD, LH and PCK have no competing interests. JSB is a founder, director, and equity holder of Neochromosome, Inc. Neochromosome is developing a yeast synthetic biology platform for pathway-level antimicrobial screens.

---

**Figure S1:**
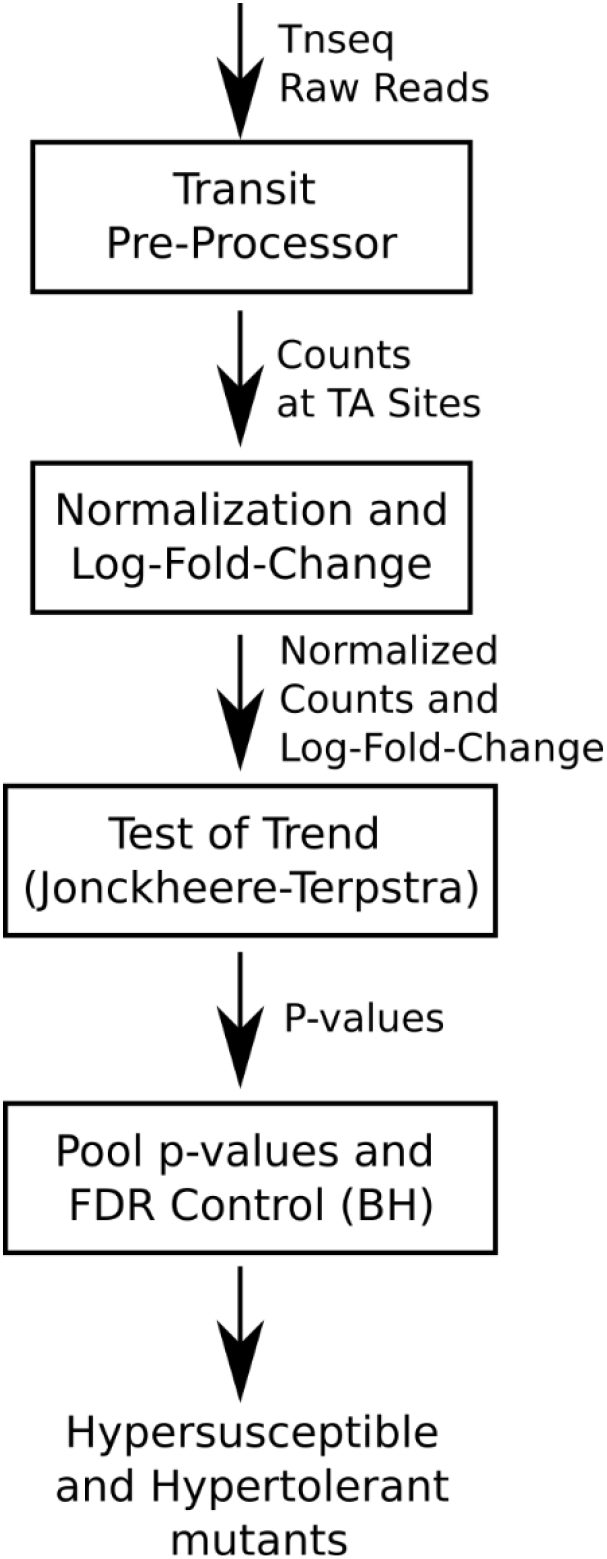
Schematic of the computational pipeline. The raw reads from Tn-seq were input into tpp, outputting a count for each TA position in the MAC109 genome. Counts were then normalized and log-fold changes calculated. A non-parametric test of trend (Jonckheere-Terpstra Test) was used to calculate p-values for each TA site. P-values for TA sites in the same gene were pooled using Stouffer’s method. Statistically significant mutants were selected based on the log-fold changes and Benjamini-Hochberg FDR-adjusted p-values. See Methods for additional details of these steps.

**Figure S2:**
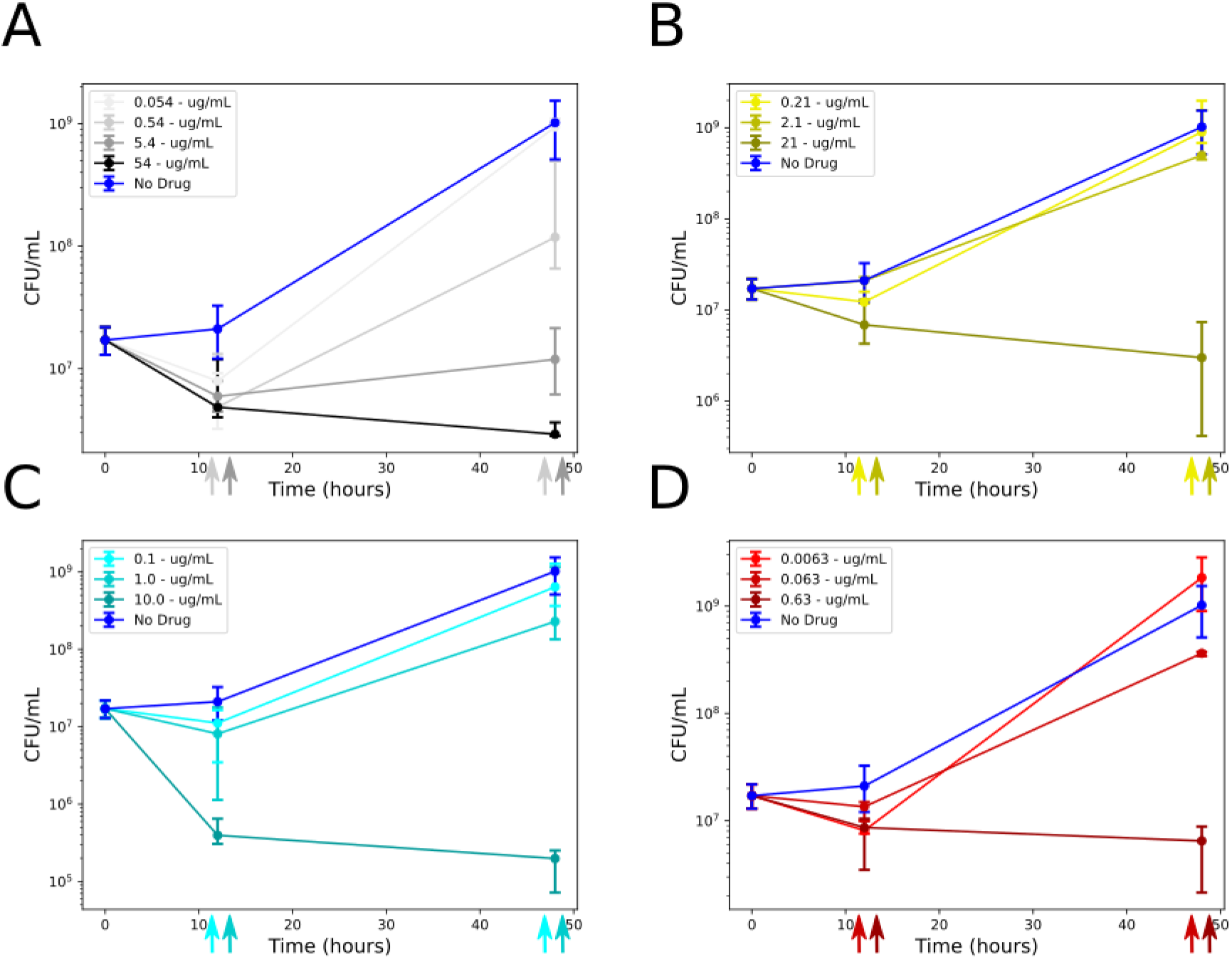
Bacterial viability of cultures by drug, dose, and time point (3 replicates each). Error bars show the minimum, maximum, and median. The arrows at the bottom indicate time point and doses used for Tn-seq. Drug-free controls (blue) are replotted in each subpanel for reference. All doses and time points were collected for drug-free controls.

**Figure S3:**
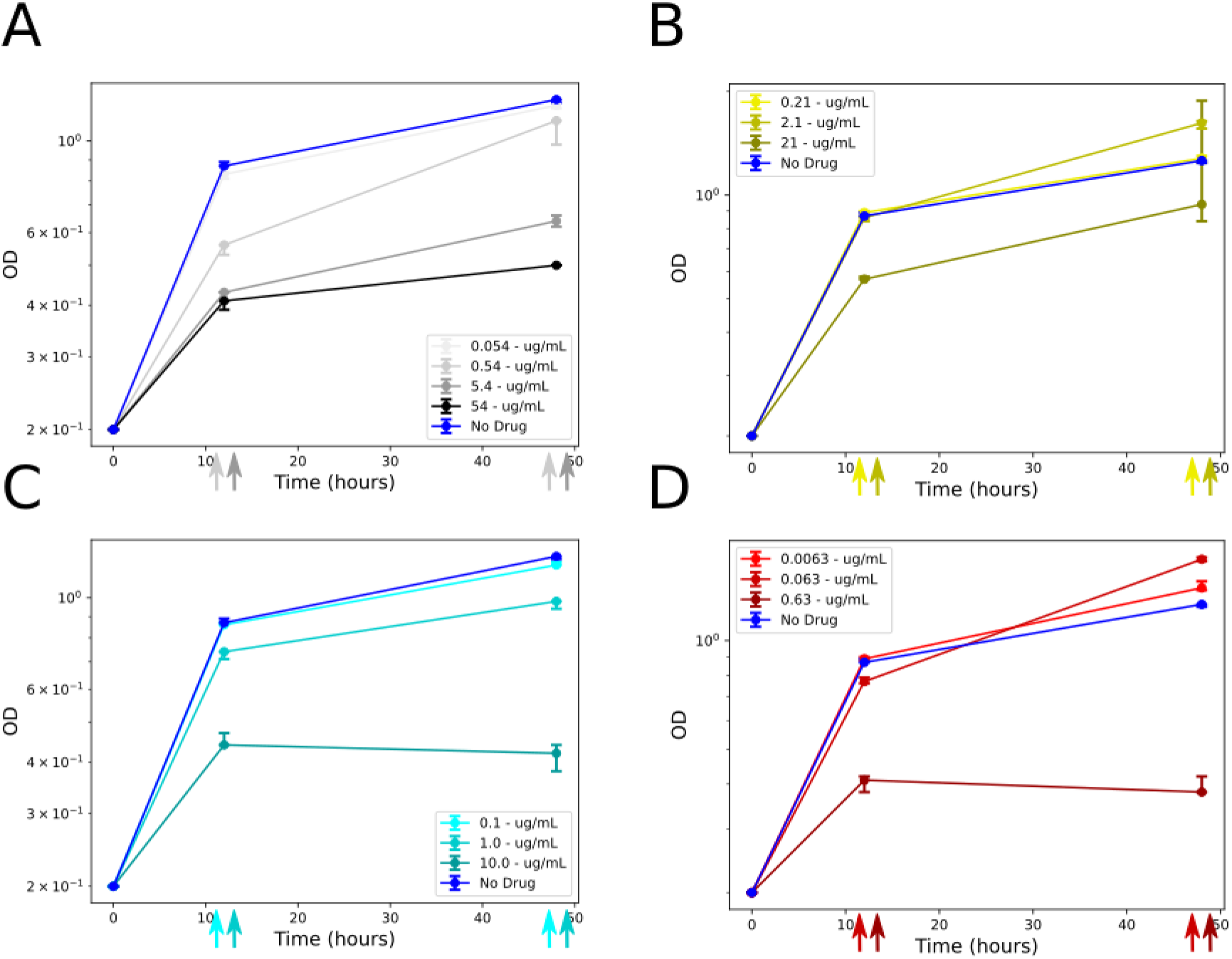
OD_600_ values by drug, dose, and time point. The arrows at the bottom indicate time point and doses used for Tn-seq. Drug-free controls (blue) are replotted in each subpanel for reference. All doses and time points were used for drug-free controls.

## Tables

***Table S1:*** Summary prediction table for antibiotic hypersusceptible MAC109 mutants.

***Table S2:*** Summary prediction table for antibiotic hypertolerant MAC109 mutants.

***Tables S3:*** Clarithromycin-hypersusceptible and hypertolerant MAC109 transposon mutants by dose and time point.

***Tables S4:*** Moxifloxacin-hypersusceptible and hypertolerant MAC109 transposon mutants by dose and time point.

***Tables S5:*** Rifabutin-hypersusceptible and hypertolerant MAC109 transposon mutants by dose and time point.

***Tables S6:*** Ethambutol-hypersusceptible and hypertolerant MAC109 transposon mutants by dose and time point.

## References

1. Falkinham JO. Epidemiology of infection by nontuberculous mycobacteria. Clin Microbiol Rev. 1996 Apr;9(2):177–215.

2. Wolinsky E. Nontuberculous mycobacteria and associated diseases. Am Rev Respir Dis. 1979 Jan;119(1):107–59.

3. Inderlied CB, Kemper CA, Bermudez LE. The Mycobacterium avium complex. Clin Microbiol Rev. 1993 Jul;6(3):266–310.

4. Ito Y, Hirai T, Maekawa K, Fujita K, Imai S, Tatsumi S, et al. Predictors of 5-year mortality in pulmonary Mycobacterium avium-intracellulare complex disease. Int J Tuberc Lung Dis. 2012;16(3):408–14.

5. Prevots DR, Shaw PA, Strickland D, Jackson LA, Raebel MA, Blosky MA, et al. Nontuberculous mycobacterial lung disease prevalence at four integrated health care delivery systems. Am J Respir Crit Care Med. 2010 Oct 1;182(7):970–6.

6. Spaulding AB, Lai YL, Zelazny AM, Olivier KN, Kadri SS, Prevots DR, et al. Geographic Distribution of Nontuberculous Mycobacterial Species Identified among Clinical Isolates in the United States, 2009-2013. Ann Am Thorac Soc. 2017 Nov;14(11): 1655–61.

7. Henkle E, Hedberg K, Schafer S, Novosad S, Winthrop KL. Population-based Incidence of Pulmonary Nontuberculous Mycobacterial Disease in Oregon 2007 to 2012. Ann Am Thorac Soc. 2015 May;12(5):642–7.

8. Winthrop KL, Marras TK, Adjemian J, Zhang H, Wang P, Zhang Q. Incidence and Prevalence of Nontuberculous Mycobacterial Lung Disease in a Large U.S. Managed Care Health Plan, 2008-2015. Ann Am Thorac Soc. 2020;17(2):178–85.

9. Hoefsloot W, van Ingen J, Andrejak C, Angeby K, Bauriaud R, Bemer P, et al. The geographic diversity of nontuberculous mycobacteria isolated from pulmonary samples: an NTM-NET collaborative study. Eur Respir J. 2013 Dec;42(6):1604–13.

10. Griffith DE, Adjemian J, Brown-Elliott BA, Philley JV, Prevots DR, Gaston C, et al. Semiquantitative Culture Analysis during Therapy for Mycobacterium avium Complex Lung Disease. Am J Respir Crit Care Med. 2015 Sep 15;192(6):754–60.

11. Daley CL, Iaccarino JM, Lange C, Cambau E, Wallace RJ, Andrejak C, et al. Treatment of Nontuberculous Mycobacterial Pulmonary Disease: An Official ATS/ERS/ESCMID/IDSA Clinical Practice Guideline. Clin Infect Dis. 2020 Aug 14;71(4):905–13.

12. Griffith DE. Therapy of nontuberculous mycobacterial disease. Curr Opin Infect Dis. 2007 Apr;20(2):198–203.

13. Parker H, Lorenc R, Ruelas Castillo J, Karakousis PC. Mechanisms of Antibiotic Tolerance in Mycobacterium avium Complex: Lessons From Related Mycobacteria. Front Microbiol. 2020;11:573983.

14. Tomasz A, Albino A, Zanati E. Multiple antibiotic resistance in a bacterium with suppressed autolytic system. Nature. 1970 Jul 11;227(5254):138–40.

15. Liao J, Sauer K. The MerR-like transcriptional regulator BrlR contributes to Pseudomonas aeruginosa biofilm tolerance. J Bacteriol. 2012 Sep;194(18):4823–36.

16. Sadovskaya I, Vinogradov E, Li J, Hachani A, Kowalska K, Filloux A. High-level antibiotic resistance in Pseudomonas aeruginosa biofilm: the ndvB gene is involved in the production of highly glycerol-phosphorylated beta-(1->3)-glucans, which bind aminoglycosides. Glycobiology. 2010 Jul;20(7):895–904.

17. Thayil SM, Morrison N, Schechter N, Rubin H, Karakousis PC. The role of the novel exopolyphosphatase MT0516 in Mycobacterium tuberculosis drug tolerance and persistence. PLoS ONE. 2011;6(11):e28076.

18. Boutte CC, Crosson S. Bacterial lifestyle shapes stringent response activation. Trends Microbiol. 2013 Apr;21(4):174–80.

19. Chuang Y-M, Belchis DA, Karakousis PC. The polyphosphate kinase gene ppk2 is required for Mycobacterium tuberculosis inorganic polyphosphate regulation and virulence. MBio. 2013 May 21;4(3):e00039–00013.

20. Chuang Y-M, Bandyopadhyay N, Rifat D, Rubin H, Bader JS, Karakousis PC. Deficiency of the novel exopolyphosphatase Rv1026/PPX2 leads to metabolic downshift and altered cell wall permeability in Mycobacterium tuberculosis. mBio. 2015 Mar 17;6(2):e02428.

21. Rodrigues L, Sampaio D, Couto I, Machado D, Kern WV, Amaral L, et al. The role of efflux pumps in macrolide resistance in Mycobacterium avium complex. Int J Antimicrob Agents. 2009 Dec;34(6):529–33.

22. Rodrigues L, Machado D, Couto I, Amaral L, Viveiros M. Contribution of efflux activity to isoniazid resistance in the Mycobacterium tuberculosis complex. Infect Genet Evol. 2012 Jun;12(4):695–700.

23. Gupta AK, Katoch VM, Chauhan DS, Sharma R, Singh M, Venkatesan K, et al. Microarray analysis of efflux pump genes in multidrug-resistant Mycobacterium tuberculosis during stress induced by common anti-tuberculous drugs. Microb Drug Resist. 2010 Mar;16(1):21–8.

24. Rojony R, Martin M, Campeau A, Wozniak JM, Gonzalez DJ, Jaiswal P, et al. Quantitative analysis of Mycobacterium avium subsp. hominissuis proteome in response to antibiotics and during exposure to different environmental conditions. Clin Proteomics. 2019;16:39.

25. Archuleta RJ, Yvonne Hoppes P, Primm TP. Mycobacterium avium enters a state of metabolic dormancy in response to starvation. Tuberculosis (Edinb). 2005 May;85(3): 147–58.

26. Sangari FJ, Parker A, Bermudez LE. Mycobacterium avium interaction with macrophages and intestinal epithelial cells. Front Biosci. 1999 Jul 15;4:D582–588.

27. McKinney JD, Höner zu Bentrup K, Muñoz-Elías EJ, Miczak A, Chen B, Chan WT, et al. Persistence of Mycobacterium tuberculosis in macrophages and mice requires the glyoxylate shunt enzyme isocitrate lyase. Nature. 2000 Aug 17;406(6797):735–8.

28. Wayne LG, Hayes LG. An in vitro model for sequential study of shiftdown of Mycobacterium tuberculosis through two stages of nonreplicating persistence. Infect Immun. 1996 Jun;64(6):2062–9.

29. Wayne LG, Lin KY. Glyoxylate metabolism and adaptation of Mycobacterium tuberculosis to survival under anaerobic conditions. Infect Immun. 1982 Sep;37(3): 1042–9.

30. van Opijnen T, Bodi KL, Camilli A. Tn-seq: high-throughput parallel sequencing for fitness and genetic interaction studies in microorganisms. Nat Methods. 2009 Oct;6(10):767–72.

31. Langridge GC, Phan M-D, Turner DJ, Perkins TT, Parts L, Haase J, et al. Simultaneous assay of every Salmonella Typhi gene using one million transposon mutants. Genome Res. 2009 Dec;19(12):2308–16.

32. Goodman AL, McNulty NP, Zhao Y, Leip D, Mitra RD, Lozupone CA, et al. Identifying genetic determinants needed to establish a human gut symbiont in its habitat. Cell Host Microbe. 2009 Sep 17;6(3):279–89.

33. Gawronski JD, Wong SMS, Giannoukos G, Ward DV, Akerley BJ. Tracking insertion mutants within libraries by deep sequencing and a genome-wide screen for Haemophilus genes required in the lung. Proc Natl Acad Sci U S A. 2009 Sep 22;106(38): 16422–7.

34. DeJesus MA, Gerrick ER, Xu W, Park SW, Long JE, Boutte CC, et al. Comprehensive Essentiality Analysis of the Mycobacterium tuberculosis Genome via Saturating Transposon Mutagenesis. MBio. 2017 Jan 17;8(1).

35. Matern WM, Jenquin RL, Bader JS, Karakousis PC. Identifying the essential genes of Mycobacterium avium subsp. hominissuis with Tn-Seq using a rank-based filter procedure. Scientific Reports. 2020 Dec;10(1):1095.

36. Xu W, DeJesus MA, Rücker N, Engelhart CA, Wright MG, Healy C, et al. Chemical genomic interaction profiling reveals determinants of intrinsic antibiotic resistance in Mycobacterium tuberculosis. Antimicrob Agents Chemother. 2017 Sep 11;

37. Gelman E, McKinney JD, Dhar N. Malachite green interferes with postantibiotic recovery of mycobacteria. Antimicrob Agents Chemother. 2012 Jul;56(7):3610–4.

38. Matern WM, Bader JS, Karakousis PC. Genome analysis of Mycobacterium avium subspecies hominissuis strain 109. Sci Data. 2018 Dec 4;5:180277.

39. Fish DN, Gotfried MH, Danziger LH, Rodvold KA. Penetration of clarithromycin into lung tissues from patients undergoing lung resection. Antimicrob Agents Chemother. 1994 Apr;38(4):876–8.

40. Blaschke TF, Skinner MH. The clinical pharmacokinetics of rifabutin. Clin Infect Dis. 1996 Apr;22 Suppl 1:S15–21; discussion S21-22.

41. Prideaux B, Via LE, Zimmerman MD, Eum S, Sarathy J, O’Brien P, et al. The association between sterilizing activity and drug distribution into tuberculosis lesions. Nat Med. 2015 Oct;21(10): 1223–7.

42. Liss RH, Letourneau RJ, Schepis JP. Distribution of ethambutol in primate tissues and cells. Am Rev Respir Dis. 1981 May;123(5):529–32.

43. van Ingen J, Egelund EF, Levin A, Totten SE, Boeree MJ, Mouton JW, et al. The pharmacokinetics and pharmacodynamics of pulmonary Mycobacterium avium complex disease treatment. Am J Respir Crit Care Med. 2012 Sep 15;186(6):559–65.

44. DeJesus MA, Ambadipudi C, Baker R, Sassetti C, Ioerger TR. TRANSIT--A Software Tool for Himar1 TnSeq Analysis. PLoS Comput Biol. 2015 Oct;11(10):e1004401.

45. Jonckheere AR. A Distribution-Free k-Sample Test Against Ordered Alternatives. Biometrika. 1954 Jun;41(1/2):133.

46. Terpstra, T.J. The Asymptotic Normality and Consistency of Kendall’s Test against Trend, When Ties Are Present in One Ranking. Indag Math. 1952;14(3):327–33.

47. Matern W. wmatern/hypersusceptibility_MAC109: Release with DOI (zenodo) [Internet]. Zenodo; 2021 [cited 2021 Feb 15]. Available from: https://zenodo.org/record/4542412

48. Braunstein M, Brown AM, Kurtz S, Jacobs WR. Two nonredundant SecA homologues function in mycobacteria. J Bacteriol. 2001 Dec;183(24):6979–90.

49. Ligon LS, Rigel NW, Romanchuk A, Jones CD, Braunstein M. Suppressor analysis reveals a role for secY in the secA2-dependent protein export pathway of mycobacteria. Journal of Bacteriology. 2013;195(19):4456–65.

50. Sullivan JT, Young EF, Mccann JR, Braunstein M. The Mycobacterium tuberculosis SecA2 system subverts phagosome maturation to promote growth in macrophages. Infection and Immunity. 2012 Mar;80(3):996–1006.

51. Zulauf KE, Sullivan JT, Braunstein M. The SecA2 pathway of Mycobacterium tuberculosis exports effectors that work in concert to arrest phagosome and autophagosome maturation. PLoS Pathogens. 2018 Apr 1;14(4).

52. Karakousis PC. Mechanisms of Action and Resistance of Antimycobacterial Agents. In: Mayers DL, editor. Antimicrobial Drug Resistance [Internet]. Totowa, NJ: Humana Press; 2009 [cited 2021 Feb 8]. p. 271–91. Available from: http://link.springer.com/10.1007/978-1-59745-180-2_24

53. Dorman CJ. DNA supercoiling and transcription in bacteria: a two-way street. BMC Mol Cell Biol. 2019 Jul 18;20(1):26.

54. Pham TDM, Ziora ZM, Blaskovich MAT. Quinolone antibiotics. Medchemcomm. 2019 Oct 1;10(10):1719–39.

55. Gupta R, Barkan D, Redelman-Sidi G, Shuman S, Glickman MS. Mycobacteria exploit three genetically distinct DNA double-strand break repair pathways. Molecular Microbiology. 2011 Jan;79(2):316–30.

56. Choudhary E, Sharma R, Kumar Y, Agarwal N. Conditional silencing by CRISPRi reveals the role of DNA gyrase in formation of drug-tolerant persister population in Mycobacterium tuberculosis. Frontiers in Cellular and Infection Microbiology. 2019;9(MAR):70.

57. Malik M, Chavda K, Zhao X, Shah N, Hussain S, Kurepina N, et al. Induction of mycobacterial resistance to quinolone class antimicrobials. Antimicrobial Agents and Chemotherapy. 2012 Jul;56(7):3879–87.

58. Wipperman MF, Heaton BE, Nautiyal A, Adefisayo O, Evans H, Gupta R, et al. Mycobacterial Mutagenesis and Drug Resistance Are Controlled by Phosphorylation- and Cardiolipin-Mediated Inhibition of the RecA Coprotease. Molecular Cell. 2018 Oct 4;72(1):152–161.e7.

59. Pisu D, Provvedi R, Espinosa DM, Payan JB, Boldrin F, Palù G, et al. The Alternative Sigma Factors SigE and SigB Are Involved in Tolerance and Persistence to Antitubercular Drugs. Antimicrobial Agents and Chemotherapy. 2017 Dec;61(12):e01596–17, e01596-17.

60. Vandal OH, Roberts JA, Odaira T, Schnappinger D, Nathan CF, Ehrt S. Acid-susceptible mutants of Mycobacterium tuberculosis share hypersusceptibility to cell wall and oxidative stress and to the host environment. J Bacteriol. 2009 Jan;191(2):625–31.

61. Nguyen D, Joshi-Datar A, Lepine F, Bauerle E, Olakanmi O, Beer K, et al. Active Starvation Responses Mediate Antibiotic Tolerance in Biofilms and Nutrient-Limited Bacteria. Science. 2011 Nov 18;334(6058):982–6.

62. Dutta NK, Klinkenberg LG, Vazquez M-J, Segura-Carro D, Colmenarejo G, Ramon F, et al. Inhibiting the stringent response blocks Mycobacterium tuberculosis entry into quiescence and reduces persistence. Sci Adv. 2019 Mar;5(3):eaav2104.

63. Ehrt S, Schnappinger D, Rhee KY. Metabolic principles of persistence and pathogenicity in Mycobacterium tuberculosis. Nat Rev Microbiol. 2018 Aug;16(8):496–507.

64. Meylan S, Porter CBM, Yang JH, Belenky P, Gutierrez A, Lobritz MA, et al. Carbon Sources Tune Antibiotic Susceptibility in Pseudomonas aeruginosa via Tricarboxylic Acid Cycle Control. Cell Chem Biol. 2017 Feb 16;24(2):195–206.

65. Zalis EA, Nuxoll AS, Manuse S, Clair G, Radlinski LC, Conlon BP, et al. Stochastic Variation in Expression of the Tricarboxylic Acid Cycle Produces Persister Cells. mBio. 2019 Sep 17;10(5).

66. Rowe SE, Wagner NJ, Li L, Beam JE, Wilkinson AD, Radlinski LC, et al. Reactive oxygen species induce antibiotic tolerance during systemic Staphylococcus aureus infection. Nat Microbiol. 2020 Feb;5(2):282–90.

67. Thomas VC, Chittezham Thomas V, Kinkead LC, Janssen A, Schaeffer CR, Woods KM, et al. A dysfunctional tricarboxylic acid cycle enhances fitness of Staphylococcus epidermidis during β-lactam stress. mBio. 2013 Aug 20;4(4).

68. Singh KS, Sharma R, Keshari D, Singh N, Singh SK. Down-regulation of malate synthase in Mycobacterium tuberculosis H37Ra leads to reduced stress tolerance, persistence and survival in macrophages. Tuberculosis. 2017 Sep;106:73–81.

69. Nandakumar M, Nathan C, Rhee KY. Isocitrate lyase mediates broad antibiotic tolerance in Mycobacterium tuberculosis. Nat Commun. 2014 Sep;5(1):4306.

70. Andréjak C, Almeida DV, Tyagi S, Converse PJ, Ammerman NC, Grosset JH. Characterization of mouse models of Mycobacterium avium complex infection and evaluation of drug combinations. Antimicrob Agents Chemother. 2015 Apr;59(4):2129–35.

71. Verma D, Stapleton M, Gadwa J, Vongtongsalee K, Schenkel AR, Chan ED, et al. Mycobacterium avium Infection in a C3HeB/FeJ Mouse Model. Front Microbiol. 2019;10:693.

